# IMPACT: Genomic annotation of cell-state-specific regulatory elements inferred from the epigenome of bound transcription factors

**DOI:** 10.1101/366864

**Authors:** Tiffany Amariuta, Yang Luo, Steven Gazal, Emma E. Davenport, Bryce van de Geijn, Harm-Jan Westra, Nikola Teslovich, Yukinori Okada, Kazuhiko Yamamoto, RACI consortium, GARNET consortium, Alkes Price, Soumya Raychaudhuri

## Abstract

Despite significant progress in annotating the genome with experimental methods, much of the regulatory noncoding genome remains poorly defined. Here we assert that regulatory elements may be characterized by leveraging local epigenomic signatures at sites where specific transcription factors (TFs) are bound. To link these two identifying features, we introduce IMPACT, a genome annotation strategy which identifies regulatory elements defined by cell-state-specific TF binding profiles, learned from 515 chromatin and sequence annotations. We validate IMPACT using multiple compelling applications. First, IMPACT predicts TF motif binding with high accuracy (average AUC 0.92, s.e. 0.03; across 8 TFs), a significant improvement (all p<6.9e-15) over intersecting motifs with open chromatin (average AUC 0.66, s.e. 0.11). Second, an IMPACT annotation trained on RNA polymerase II is more enriched for peripheral blood cis-eQTL variation (N=3,754) than sequence based annotations, such as promoters and regions around the TSS, (permutation p<1e-3, 25% average increase in enrichment). Third, integration with rheumatoid arthritis (RA) summary statistics from European (N=38,242) and East Asian (N=22,515) populations revealed that the top 5% of CD4+ Treg IMPACT regulatory elements capture 85.7% (s.e. 19.4%) of RA h2 (p<1.6e-5) and that the top 9.8% of Treg IMPACT regulatory elements, consisting of all SNPs with a non-zero annotation value, capture 97.3% (s.e. 18.2%) of RA h2 (p<7.6e-7), the most comprehensive explanation for RA h2 to date. In comparison, the average RA h2 captured by compared CD4+ T histone marks is 42.3% and by CD4+ T specifically expressed gene sets is 36.4%. Finally, integration with RA fine-mapping data (N=27,345) revealed a significant enrichment (2.87, p<8.6e-3) of putatively causal variants across 20 RA associated loci in the top 1% of CD4+ Treg IMPACT regulatory regions. Overall, we find that IMPACT generalizes well to other cell types in identifying complex trait associated regulatory elements.

---

Transcriptional regulation is the foundation for many complex biological phenotypes, from gene expression to disease susceptibility. However, the complexity of gene regulation, controlled by more than 1,600 human transcription factors (TFs)^1^ influencing some 20,000 protein coding gene promoters, has made functional annotation of the regulome difficult. Tens of thousands of genomic annotations have been experimentally generated, enabling the success of unsupervised methods such as chromHMM^2^ and Segway^3^ to identify global chromatin patterns that better characterize genomic function. However, linking specific regulatory processes to these identified patterns is challenging. Furthermore, although genome-wide association studies (GWAS) have identified ~10,000 trait associated variants across hundreds of polygenic traits^4^, most variants lie in noncoding regulatory regions with uncertain function.

With continually increasing numbers of genomic annotations generated from high-throughput experimental assays, *in-silico* functional characterization of variants has growing potential. These assays include genome-wide open chromatin, histone mark, and RNA expression profiling, each separately possible at the single cell level. Initially contributed by genomic consortia, such as ENCODE^5^ and Roadmap^6^, these assays have become more common place as easy-to-implement protocols have been developed, thereby contributing to the growing rate of genomic annotation generation.

Recently, integration of datasets, particularly those indicating regulatory elements, with GWAS data has successfully led to the identification of categories of disease-driving variants enriched for genetic heritability (h2)^7–9^. Such regulatory annotations identify active promoters and enhancers through open chromatin or histone mark occupancy assays in a cell type of interest^7,8,10–13^. However, these annotations include both cell-type-specific and nonspecific elements, the latter of which may affect a wide range of cellular functions that are not necessarily intrinsic to disease-driving cell-states.

Therefore, we hypothesized that the identification of regulatory elements specifically driving functional states would help us to not only better characterize regulatory elements genome-wide, but also better capture polygenic h2 of complex traits and diseases. Once the most enriched classes of regulatory elements are recognized, then it may become possible to generate biologically-founded mechanistic hypotheses.

Here, we introduce IMPACT (Inference and Modeling of Phenotype-related ACtive Transcription), a diversely applicable genome annotation strategy to predict cell-state-specific regulatory elements. We take a two-step approach to define IMPACT regulatory elements. First, we choose a single key TF, known to regulate a cell-state-specific process, and then identify binding motifs genome-wide, distinguishing between those that are bound and unbound using genomic occupancy identified by ChIP-seq in the corresponding cell-state. Second, IMPACT predicts TF occupancy at binding motifs by aggregating and performing feature selection on 503 cell-type-specific epigenomic features and 12 sequence features in an elastic net logistic regression model (**Methods**: *IMPACT Model*). Epigenomic features include histone mark ChIP-seq, ATAC-seq, DNase-seq and HiChIP (**Table S1**) assayed in hematopoietic, adrenal, brain, cardiovascular, gastrointestinal, skeletal, and other cell types, while sequence features include coding, intergenic, etc. The IMPACT model framework can easily be expanded to accommodate thousands of epigenomic feature annotations and is amenable to the increasing rates of data generation. From this regression we learn a TF binding chromatin profile, which IMPACT uses to probabilistically annotate the genome at nucleotide-resolution. We refer to high scoring regions as cell-state-specific regulatory elements (**Figure 1A**).

**Figure 1.**
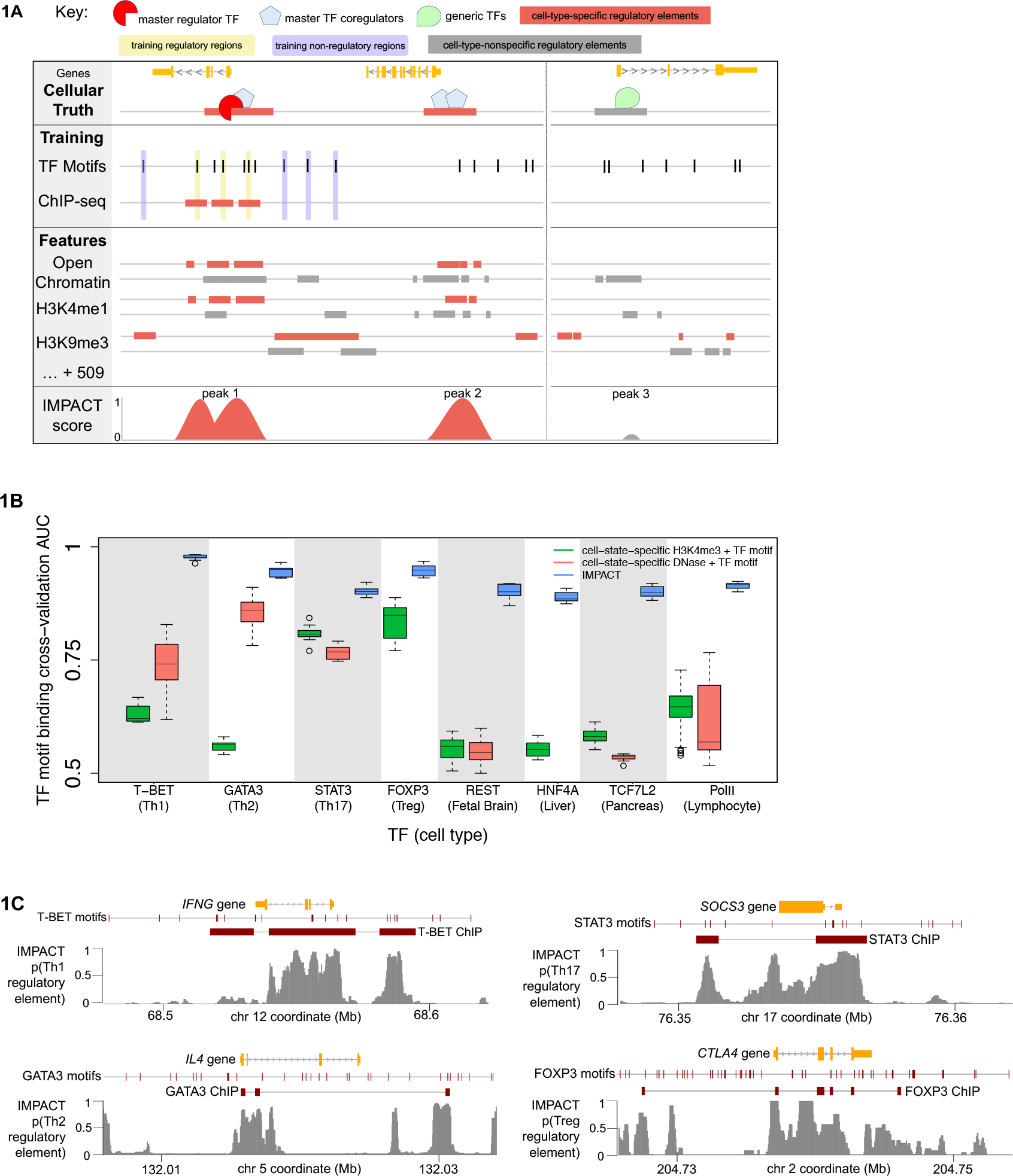
**(a)** IMPACT learns a chromatin profile of cell-type-specific regulation, characteristic of the master regulator TF (red) “gold standard” regulatory elements (TF motif and ChIP seq peak, yellow) and non-regulatory elements (TF motif, no ChIP-seq peak, purple). In this toy example, IMPACT learns that cell-type-specific open chromatin and H3K4me1 are strong predictors of cell-type-specific regulatory elements, while cell-type-nonspecific DHS and H3K4me1 are less informative. IMPACT also learns that H3K9me3 is a strong predictor of non-regulatory elements. IMPACT is expected to re-identify regulatory elements marked by master TF binding (ChIP-seq and motif) [peak 1], while also identifying others where the chromatin profile is similar, perhaps representing cell-type-specific transcriptional processes [peak 2]. IMPACT is not expected to predict regulation at cell-type-nonspecific elements [peak 3], such as promoters of generic housekeeping genes, assuming these elements have different chromatin profiles. **(b)** IMPACT significantly outperforms cell-state-specific active promoter (H3K4me3, green) and open chromatin (DNase, red) annotations in predicting TF binding on a motif over 10 trials measured by computing the average ROC AUC (receiver operator characteristic area under the curve). As there is no Treg DNase data, there is no comparison AUC distribution. **(c)** Cell-state-specific regulatory element IMPACT predictions for canonical target genes of T-BET, GATA3, STAT3, and FOXP3.

The IMPACT model assumes that binding sites of a given TF may be characterized by a consensus epigenomic signature, which may also characterize binding sites of other related TFs. If this signature is robust, each TF binding site genome-wide should share some of its characteristics. Therefore, IMPACT should be able to predict with high accuracy genome-wide TF occupancy of a motif, which has been challenging^14^. To test this model assumption, we used IMPACT to predict regulatory elements based on experimental binding identified via ChIP-seq of eight TFs assayed in eight different cell-states: T-BET, GATA3, STAT3, FOXP3, REST, HNF4A, TCF7L2, and RNA Polymerase (Pol) II in CD4+ Th(T helper)1, CD4+ Th2, CD4+ Th17, CD4+ Treg (T regulatory), fetal brain, liver, pancreatic, and lymphocytic cells, respectively (**Table S2**)^5,15–20^ (**Methods**: *IMPACT Model*). IMPACT predicts TF occupancy at binding motifs with high accuracy (average AUC across 8 TFs 0.92 (s.e. 0.03), 10-fold cross validation on 80% of data, AUC evaluated on the withheld 20%, **Figure 1B**, **Figure S1**, **Table S3**). In comparison, IMPACT significantly outperforms an approach that does not use data aggregation or a machine learning framework to predict TF binding (all *p*<6.9e-15); this approach predicts binding on motifs that overlap cell-type-specific active promoters (H3K4me3 ChIP-seq) or cell-type-specific open chromatin (DNase-seq) (average AUC 0.66 (s.e. 0.11), **Figure 1B**). We anecdotally observe that IMPACT predictions at canonical CD4+ T cell TF-specific targets (**Figure 1C**, **Figures S2-4**) substantially vary within a single ChIP-seq peak, suggesting that chromatin signatures exist on a much smaller scale and that IMPACT may be more sensitive as a regulatory annotation than TF ChIP-seq.

We developed IMPACT to model regulation specific to a functional cell-state, the most general of which may be active cellular transcription. Expression quantitative trait loci (eQTLs) are genetic variations that modulate transcription^21^. Most cis eQTLs map to TSS (transcription start site) and promoter annotations, and more rarely to the 5’ UTR (untranslated region)^22^. We hypothesized that an IMPACT annotation tracking active transcription, trained on RNA Polymerase (Pol) II binding sites, would capture cis eQTL causal variation better than the most strongly enriched canonical annotations.

We obtained SNP-level summary statistics from a large and previously published eQTL analysis on 3,754 peripheral blood samples^23^. We then used IMPACT to annotate SNPs tested in the eQTL analysis with RNA Pol II specific regulatory element probabilities, learned from Pol II binding profiles in T cells and B cells, the predominant cell populations of peripheral blood. Next, we computed a genome-wide enrichment (**Methods**: *eQTL enrichment*) of chi-squared cis eQTL association statistics, averaged over all genes, across Pol II IMPACT and several sequenced-based annotations, such as TSS windows, promoters, and enhancers, separately, revealing that Pol II IMPACT has the strongest chi-squared enrichment (1.7, permutation p<1e-3, 25% average increase in enrichment) (**Figure 2**). These results argue that Pol II IMPACT regions better localize active promoter and proximal regulatory regions driving eQTLs than the compared canonical genomic annotations, which may be less specific due to their larger sizes and restrictive binary characterization. This suggests that IMPACT may be more effective at prioritizing causal SNP variation when fine-mapping eQTLs. These results also argue that the biological basis of eQTLs are related to Pol II regions, which is a refinement over previous observations that eQTL causal variation is concentrated near and around TSS and promoter regions.

**Figure 2.**
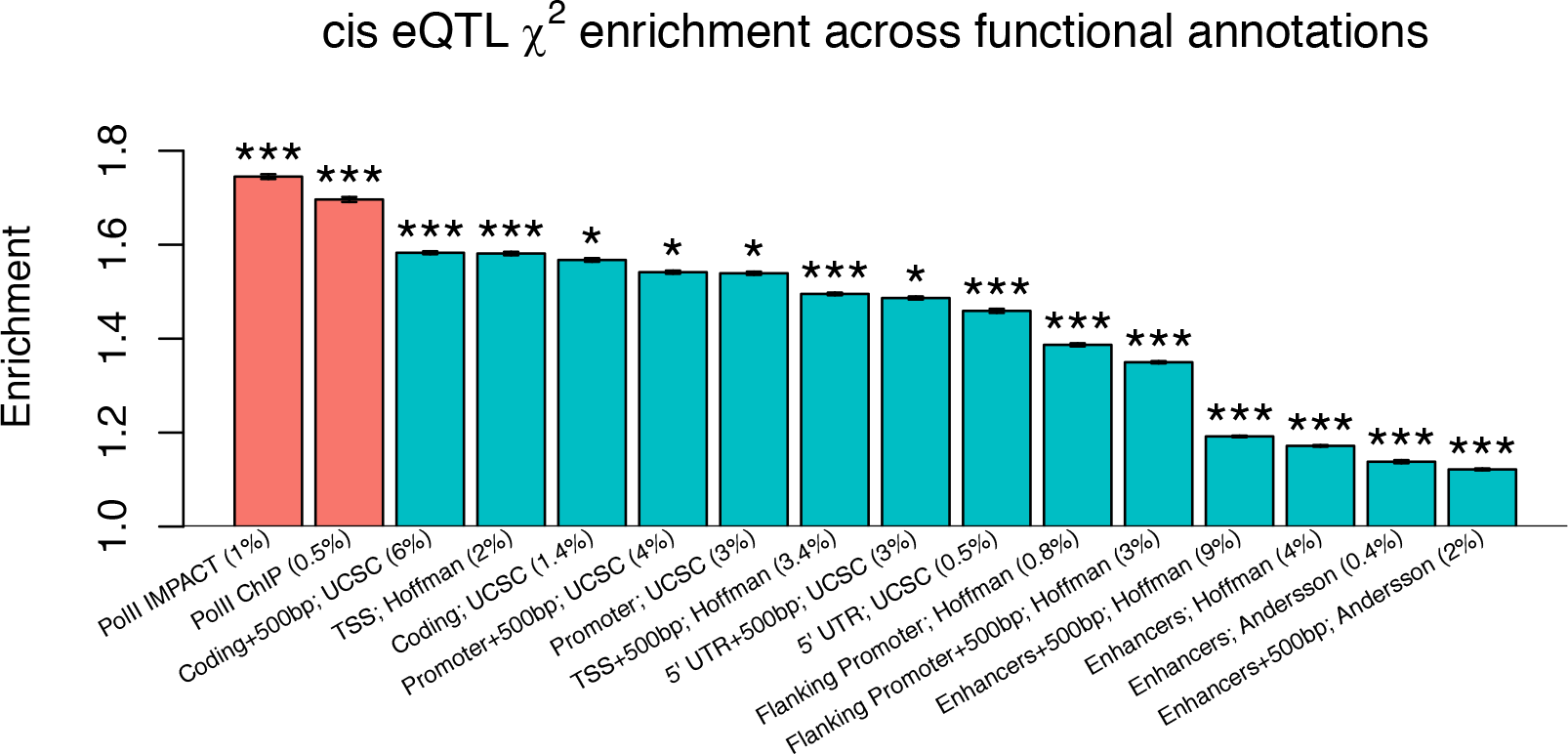
Enrichment of cis eQTL chi-squared statistics, measured in 3,754 peripheral blood samples, across functional annotations including Pol II IMPACT, Pol II ChIP, used to train IMPACT, and various sequence annotations. Values in parentheses after annotation name are the average annotation value across all common variants and represent the effective genome-wide size of the annotation. No asterisk denotes permutation *p*>0.05, 1 asterisk permutation *p*<0.05, 2 asterisks permutation *p*<0.01, 3 asterisks permutation *p*<0.001. Intervals at the top of each bar represent the 95% confidence interval of the enrichment estimate, which was averaged over 7,025 unique genes genome-wide.

We previously hypothesized that IMPACT annotations of pathogenic cell-states would more precisely capture polygenic trait h2, compared to regulatory annotations that don’t resolve cell-states. Testing this hypothesis requires a polygenic trait with a well-studied disease-driving cell type. Genetic studies of rheumatoid arthritis (RA), an autoimmune disease that attacks synovial joint tissue leading to permanent joint damage and disability^24^, have suggested a critical role by CD4+ T cells^7,8,11,12,25–29^. However, CD4+ T cells are extremely heterogeneous: naive CD4+ T cells may differentiate into memory T cells, and then into effector T cells including Th1, Th2, and Th17 and T regulatory cells, requiring the action of a limited number of key transcription factors (TFs): T-BET or STAT4, GATA3 or STAT6, STAT3 or RORγt, FOXP3 or STAT5, respectively^30^. As these CD4+ T effector cell-states contribute to RA risk^7,11,28^, we hypothesized that CD4+ T cell-state-specific IMPACT regulatory element annotations would better capture RA h2 than annotations that generalize CD4+ T cells and ignore the differential functionality of effector cell-states. Therefore, we built IMPACT annotations in four CD4+ T cell-states, Th1, Th2, Th17, and Treg. We then integrated S-LDSC (stratified linkage disequilibrium (LD) score regression)^7^ with publicly available European (EUR, N = 38,242)^7,31^ and East Asian (EAS, N = 22,515)^32^ RA GWAS summary statistics to partition the common SNP h2 of RA. While partitioning h2 with IMPACT annotations, we use a set of 69 baseline annotations, a subset of the 75 annotations referred to as the baseline-LD model^9^, specifically excluding T cell-related annotations, to control for cell-type-nonspecific functional, LD-related, and minor allele frequency (MAF) associations. We use two metrics to assess the contribution of our IMPACT annotation to RA h2: enrichment and per-annotation standardized effect size, τ*. Enrichment is defined as the proportion of h2 divided by the genome-wide proportion of SNPs in the annotation, and τ* is defined as the proportionate change in per-SNP h2 associated with a one standard deviation increase in the value of the annotation^9^. τ* captures the unique contribution of an annotation to the h2 model, conditional on other present annotations. The τ* of an annotation that successfully captures trait h2 will be significantly greater than zero; the greater the τ*, the more h2 that the annotation captures.

We found that each CD4+ T cell-state-specific IMPACT annotation is significantly enriched with RA h2 in both European and East Asian populations (average enrichment = 20.05, all *p*<1.9e-04, **Figure 3A**, **Table S4**). We estimated total genome-wide polygenic RA h2 to be about 18% for EUR and 21% for EAS (**Methods**: *S-LDSC*). As RA is primarily driven by CD4+ T cells, we expectedly find that τ* is significantly positive for all CD4+ T IMPACT annotations separately conditioned on the cell-type-nonspecific baselineLD annotations (all *p*<2.1e-03, **Figure 3B**, **Figure S5**). We then selected the top 5% of regulatory SNPs according to each CD4+ T IMPACT annotation and find that the Treg annotation explains the greatest proportion of RA h2, 85.7% (s.e. 19.4%, enrichment p<1.6e-5) meta-analyzed between both EUR and EAS populations (**Figure 3C**). Other CD4+ T cell-state annotations explain similarly large proportions of RA h2. Furthermore, we observe that the top 9.8% of CD4+ Treg IMPACT regulatory elements, consisting of all SNPs with a non-zero annotation value, capture 97.3% (s.e. 18.2%, enrichment p<7.6e-7) of RA h2 in EUR. This powerful result is the most comprehensive explanation for RA h2, to our knowledge, to date.

We then compared the enrichments and τ* of each CD4+ T IMPACT annotation to that of various T cell functional annotations, using the EUR RA summary statistics: TF binding motifs, genome-wide TF occupancy (ChIP-seq), an annotation that assigns each SNP a value proportional to the number of IMPACT epigenomic features it overlaps (Averaged Tracks), the five largest τ* CD4+ T cell-specific histone mark annotations^7^, the five largest τ* CD4+ T cell-specifically expressed gene sets and their regulatory elements^8^, and CD4+ T cell super enhancers^33^. While the CD4+ Treg IMPACT annotation has similar enrichment to CD4+ T histone marks (22.9x compared to 23.4x on average, respectively), we observe that the Treg IMPACT annotation is substantially more enriched than CD4+ T specifically expressed gene sets (22.9x compared to 2.9x on average, respectively) (**Figure 4A**, **Figure S6**). We note that the average RA h2 captured by these CD4+ T histone marks, ranging in size from 1-3% of SNPs, is 42.3%. The average RA h2 captured by these CD4+ T specifically expressed gene sets, ranging in size from 11-13% of SNPs, is 36.4%. This is a large contrast to the 85.7% of RA h2 captured by the top 5% of SNPs in the Treg IMPACT annotation. Then, we computed the annotation τ* while pairwise conditioned on each other and on the baseline-LD. We found that the CD4+ Treg and Th2 IMPACT τ* are significantly positive (all Treg τ* > 1.9, all Th2 τ* > 1.7; all Treg τ* *p*<5.0e-3, all Th2 τ* p<0.01) and more significant, thereby capturing RA h2 better, than all other T cell annotations, except H3K27ac in Th2 cells (**Figure 4B**, **Figure S6**). Previous work has reported that the threshold for impactful values of |τ*| is approximately 0.24^9^. Overall, these results suggest that IMPACT is identifying areas of concentrated RA h2 that other T cell regulatory annotations have not.

**Figure 3.**
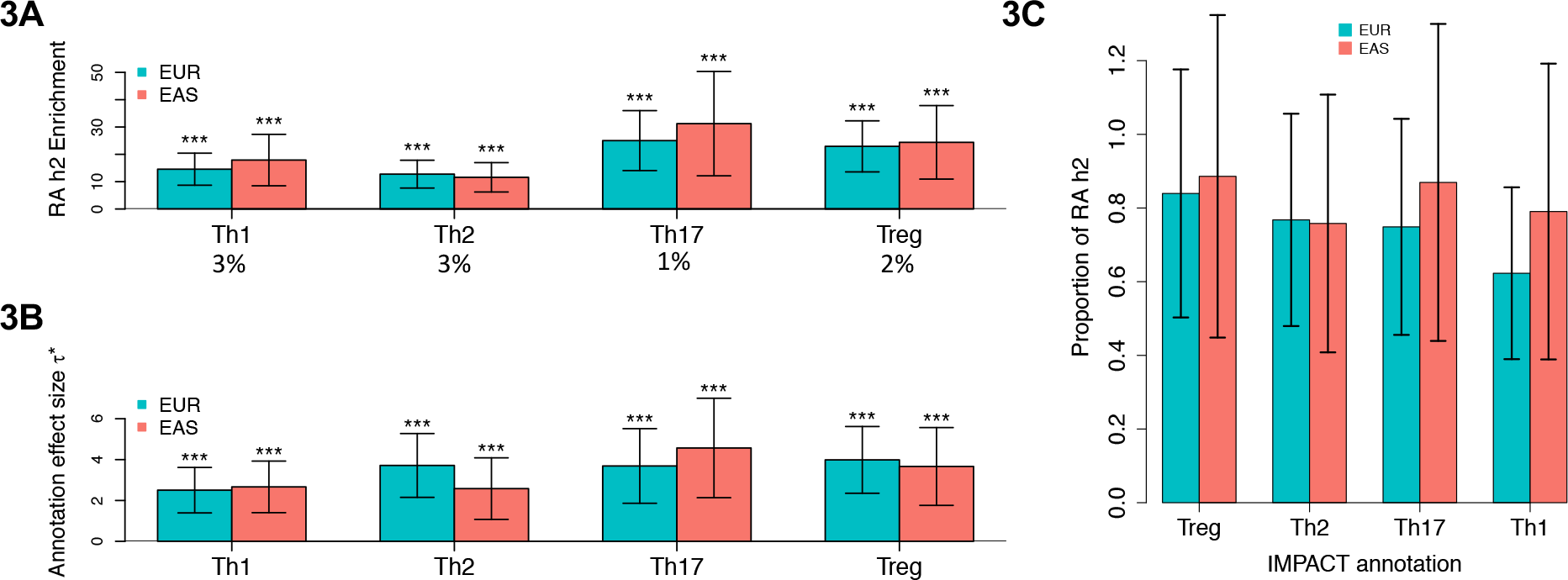
**(a)** Enrichment of RA h2 in CD4+ T IMPACT for EUR and EAS populations. Values below cell-states are the average annotation value across all common variants and represent the effective genome-wide size of the annotation. **(b)** Annotation effect size (τ*) of each annotation separately conditioned on the baseline-LD. **(c)** Proportion of total causal RA h2 explained by the top 5% of SNPs in each IMPACT annotation. For all panels, 95% CI represented by black lines. For panels a and b, no asterisk denotes *p*>0.05, 1 asterisk *p*<0.05, 2 asterisks *p*<0.01, 3 asterisks *p*<0.001.

**Figure 4.**
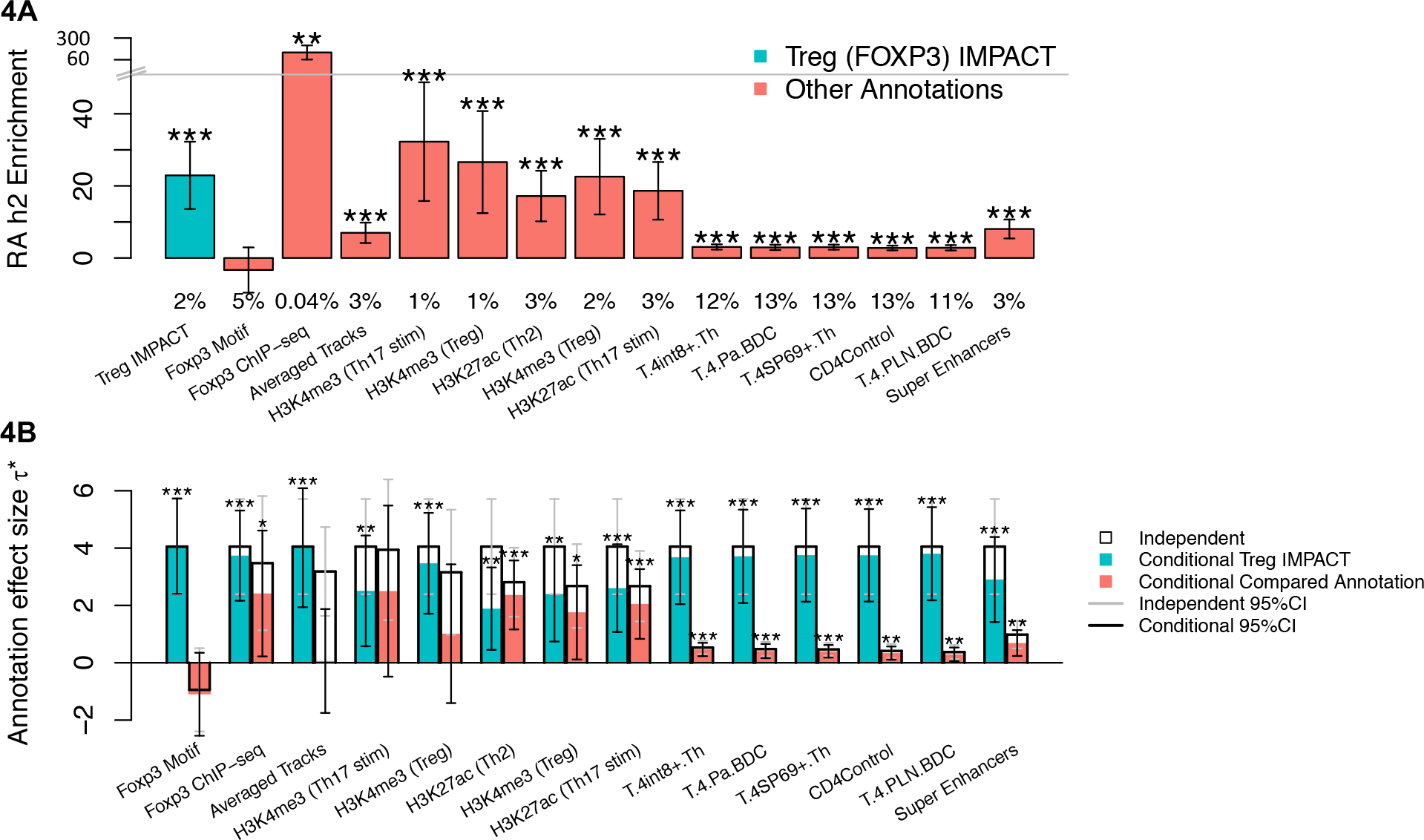
**(a)** RA h2 enrichment of CD4+ Treg related annotations and compared T cell functional annotations. Values below cell-states represent the effective genome-wide size of the annotation. From left to right, we compare Treg IMPACT to genome-wide FOXP3 motifs, FOXP3 ChIP-seq, a genome-wide averaged track of features in the IMPACT framework (Averaged Tracks), the top 5, in terms of independent τ*, cell-type-specific histone modification annotations^7^, the top 5, in terms of independent τ*, cell-type-specifically expressed gene sets (URLS)^8^, and T cell super enhancers^33^. **(b)** CD4+ Treg IMPACT annotation standardized effect size (τ*) consistently significantly greater than zero when conditioned on other T cell related functional annotations. τ* for independent (e.g. non-conditional) analyses are denoted by the top of each black bar, as a reference for the conditional analyses, denoted by the top of each colored bar. The Treg IMPACT annotation captures a significant amount of RA h2, denoted by significantly positive τ*, regardless of the conditioned annotation. For panels a and b, no asterisk denotes *p*>0.05, 1 asterisk *p*<0.05, 2 asterisks *p*<0.01, 3 asterisks *p*<0.001.

We next applied our CD4+ T IMPACT annotations to 41 additional polygenic traits^9,34,35^ and observed consistently significantly positive τ* for immune-mediated traits, such as Crohn’s, “all autoimmune disease”, respiratory ear/nose/throat, and “allergy and eczema” (mean τ* = 3.2; all *p*<5.9e-4, *p*<1.9e-5, *p*<3.6e-3, *p*<1.7e-3, respectively), and several blood traits, eosinophil and white blood cell counts (mean τ* = 2.5; all *p*<1.6e-11, p<0.02, respectively), but not for non-immune-mediated traits (**Figure 5**, **Table S5**). For comparison, we created several other IMPACT annotations in different cell types targeting h2 in a range of traits and highlight a few examples. For a liver IMPACT annotation, trained on HNF4A^5^ (hepatocyte nuclear factor 4A), τ* is positive for LDL and HDL (mean τ* = 2.0; *p*<0.02, p<1.2e-3, respectively), liver-associated traits^20,36^. For a macrophage IMPACT annotation, trained on IRF5^37^, τ* is positive for some immune-mediated and blood traits (mean τ* = 2.8, all p<8.2e-3) and intriguingly also for schizophrenia (τ* = 0.9, p<4.9e-5), as the literature supports a putative MHC association^38^. Finally, for a CD4+ Treg IMPACT annotation, trained on STAT5^19^, an alternative key TF for Tregs, the values of τ* across all traits resemble that of FOXP3. This argues that the ability of IMPACT to capture polygenic h2 does not depend on the choice of key regulator. Overall, this suggests that IMPACT is a promising strategy to identify complex trait associated regulatory elements across a range of cell-states.

**Figure 5.**
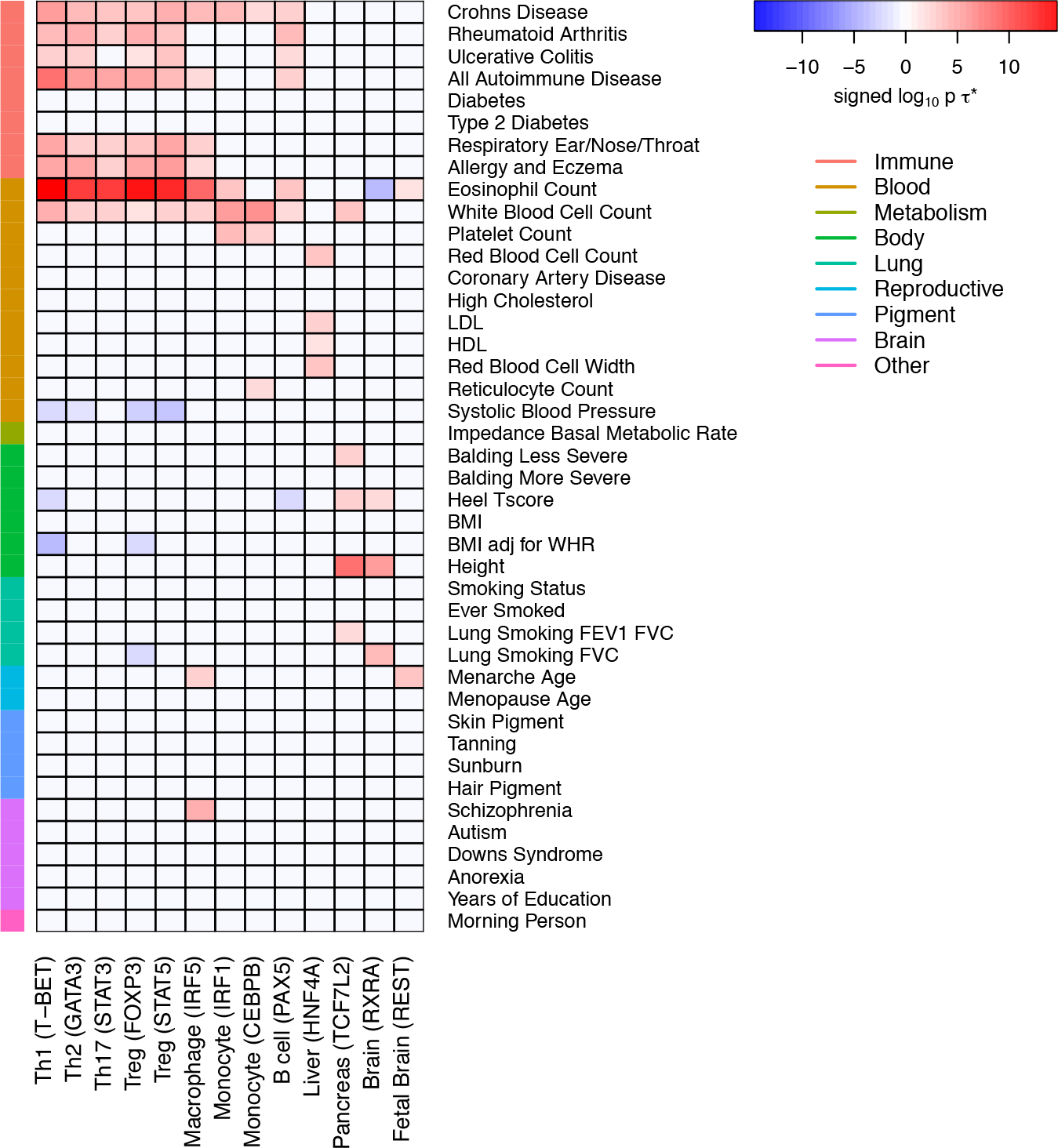
Signed log_10_ p-values of τ* for 42 traits across the four CD4+ T IMPACT and various other cell types for comparison. Annotations capture h2 in distinct sets of complex traits, shown by significantly positive τ*. Color shown only if p-value of τ* < 0.025 after multiple hypothesis correction.

As we have demonstrated the ability of IMPACT to annotate the noncoding genome, validated by captured polygenic h2, we hypothesized that this improved genomic annotation might inform functional variant fine-mapping. Using a GWAS of 11,475 European RA cases and 15,870 controls^39^, an independent study from the European summary statistics used in our h2 analyses, our group recently fine-mapped a subset of 20 RA risk loci, each with a manageable number of putatively causal variants, and created 90% credible sets of these SNPs^40^. We computed the enrichment of fine-mapped causal probabilities across these 20 loci in the top 1% of our CD4+ T cell-state-specific IMPACT annotations (**Methods**: *Posterior Probability Enrichment*). We found that the Treg annotation is significantly enriched (2.87, permutation *p*<8.6e-03, **Figure 6A**, **Table S6**), while other annotations are not. The Treg IMPACT annotation may thus be useful to prune putatively causal RA variants. Furthermore, we observe uniquely high Treg enrichment in the *BACH2* and *IRF5* loci (16.2 and 8.1, respectively), suggesting putatively causal SNPs in these loci may function in a Treg-specific context.

**Figure 6.**
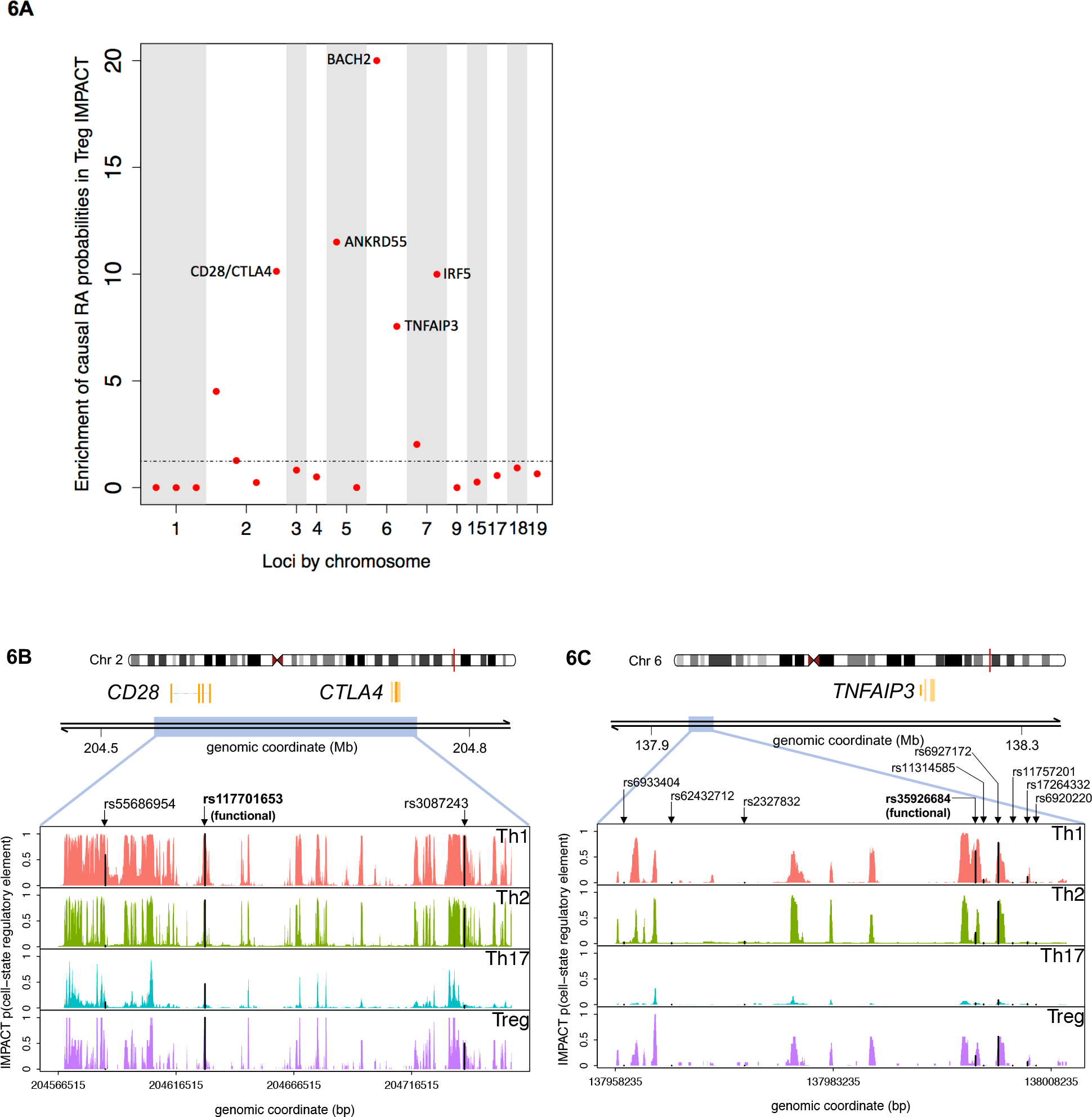
**(a)** Significant enrichment of posterior probabilities of putatively causal RA SNPs in the top 1% of SNPs annotated with CD4+ Treg regulatory element probabilities, highlighting particularly strong enrichment at the *BACH2*, *ANKRD55*, *CTLA4/CD28*, *IRF5*, and *TNFAIP3* loci. **(b,c)** IMPACT corroborates experimental validations of putatively causal RA SNPs. For two RA-associated loci, *CTLA4/CD28* and *TNFAIP3*, we examine the putatively causal SNP with experimentally validated differential enhancer activity (bolded) and other 90% credible set SNPs (unbolded)^40^. IMPACT scores at these SNPs are highlighted with a black line. **(b)** We observe high probability IMPACT regulatory elements in all four CD4+ T cell-states for the functional SNP rs117701653 in the *CD28/CTLA4* locus. **(c)** We observe a high probability Th1-specific IMPACT regulatory element for the functional SNP rs35926684 in the *TNFAIP3* locus.

In the same study, our group observed both differential binding of CD4+ T nuclear extract via EMSA and differential enhancer activity via luciferase assays at two credible set SNPs, narrowing down the list of putatively causal variants in the *CD28/CTLA4* and *TNFAIP3* loci^40^. We observed that both variants with functional activity were located at predicted IMPACT regulatory elements, suggesting that IMPACT may be used to narrow down credible sets to reduce the amount of experimental follow up. First, at the *CD28/CTLA4* locus, IMPACT predicts high probability regulatory elements across the four CD4+ T cell-states at the functional SNP rs117701653 and lower probability regulatory elements at other credible set SNPs rs55686954 and rs3087243 (**Figure 6B**). Second, at the *TNFAIP3* locus, we observe high probability regulatory elements at the functional SNP rs35926684 and other credible set SNP rs6927172 (**Figure 6C**) and do not predict regulatory elements at the other 7 credible set SNPs. The CD4+ Th1 specific regulatory element at rs35926684 suggests that this SNP may alter gene regulation specifically in Th1 cells and hence, we suggest any functional follow-up be done in this cell-state. Fewer than 11% of the credible set SNPs in the other 18 fine-mapped loci have high IMPACT cell-state-specific regulatory element probabilities (**Figure S7**). We note that disease-relevant IMPACT functional annotations may be integrated with existing functional fine mapping methods, like PAINTOR^41^ or CAVIARBF^42^, to assign causal posterior probabilities to variants.

In summary, IMPACT predicts cell-state-specific regulatory elements based on epigenomic and sequence profiles of experimental cell-state-specific TF binding. First, we observed that the robust epigenomic footprint of TF binding sites allows for accurate motif binding prediction. Second, we show the versatility of IMPACT to model functional cell-state regulatory annotations to more precisely capture causal variation in gene expression and disease susceptibility. Lastly, we demonstrated that IMPACT may identify functional variants a priori. We recognize several important limitations to our work: 1) we have not experimentally validated the activity of any of our predicted regulatory elements, 2) predicted regulatory elements are limited to genomic regions that have been epigenetically assayed, 3) IMPACT, as presented, is limited to cell-states in which ChIP-seq of a key TF has been performed. In light of these limitations, IMPACT is an emerging strategy for identifying trait associated regulatory elements and generating hypotheses about the cell-states in which variants may be functional, motivating the need to develop therapeutics that target specific disease-driving cell-states.

## Methods

### Data

#### Genome-wide Annotation Data

We obtained publicly available genome-wide annotations in a broad range of cell types for the GRCh37 (hg19) assembly. The accession numbers and or file names for features downloaded from NCBI, Blueprint, Roadmap, and chromHMM are listed in **Table S1**. Features from Finucane et al 2015^7^ are labeled as they were in supplemental tables of this study. Cell-type-specific annotation types include ATAC-seq, DNase-seq, FAIRE-seq, HiChIP, polymerase and elongation factor ChIP-seq, and histone modification ChIP-seq. Sequence annotations were downloaded from UCSC’s publicly available bedfiles and include conservation, exons, introns, intergenic regions, 3’UTR, 5’UTR, promoter-TSS, TTS, and CpG islands). All genome-wide feature data, except conservation, is represented in standard 6-column bedfile format. Conservation is represented in bedgraph format, in which the average score is reported for each 1024 bp window genome-wide.

#### TF ChIP-seq data

We determined genome-wide TF occupancy from publicly available ChIP-seq (**Table S2**) of 14 key regulators (T-BET^15,16^, GATA3^17^, STAT3^18^, FOXP3^19^, STAT5^19^, IRF5^37^, IRF1^43^, CEBPB^44^, PAX5^45^, REST^5^, RXRA^5^, HNF4A^20^, TCF7L2^5^, RNA Pol II^5,16,46^) assayed in their respective primary cell-states Th1, Th2, Th17, Tregs, Tregs, monocytes, monocytes, monocytes, B cells, fetal brain cells, brain cells, liver cells, pancreatic cells, and lymphocytes. ChIP-seq peaks were called by macs^47^ [v1.4.2 20120305] (all p<1e-5).

### IMPACT Model

We build a model that predicts TF binding on a motif by learning the epigenomic profiles of the TF binding sites. We use logistic regression to model the log odds of TF binding based on a linear combination of the effects *β*_*j*_ of the *j* epigenomic features, where *β*_0_ is an intercept.

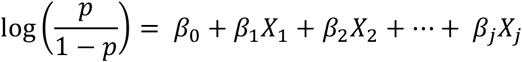

From the log odds, which ranges from negative to positive infinity, we compute the probability of TF binding, ranging from 0 to 1.

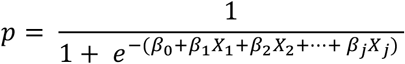

We use an elastic net logistic regression framework implemented by the *cv.glmnet* R [v1.0.143] package^48^, in which optimal *β* are fit according to the following objective function,

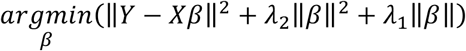

We use a regularization strategy to constrain the values of *β*and help prevent overfitting. Elastic net regularization is a compromise between the lasso (L1) and ridge (L2) penalties. Both penalties have advantageous effects on the model fit of *j* features. Lasso performs feature selection, allowing only a subset to be used in the final model, pushing some *β*_*j*_ to 0, which is useful for large feature sets such as IMPACT and avoids overfitting. When used alone, lasso will arbitrarily select one of several correlated features to be included in the final model; incorporating the ridge term limits this effect of lasso. Furthermore, ridge provides a quadratic term, making the optimization problem convex with a single optimal solution. The elastic net logistic regression has the following parameters: *alpha*, *lambda*, and *type*. Alpha is the mix term between the lasso and ridge penalties in the objective function, which controls the sparsity of betas. We set *alpha* to 0.5, such that the L1 and L2 terms equally contribute, correlated features may be retained to some degree, and not too many betas are pushed to 0. We set *lambda* to *lambda.min*, the value of *λ* that yields minimum mean cross-validated error, and *type* set to response, such that our predictions are in probability space rather than log odds space. We choose to not employ a deep learning approach in order to retain interpretability of our model *β*. More precisely, knowledge of which genomic annotations are most informative for predicting transcriptional regulation will be useful for deciding which high-throughput experimental assays should be preferentially performed in an effort to learn more about the regulome.

#### Training IMPACT

We train IMPACT on gold standard regulatory and non-regulatory elements of a particular TF, meaning that there is one IMPACT model per TF/cell-state pair. To build the regulatory class, we scanned the TF ChIP-seq peaks, mentioned above, for matches to the TF-specific binding motif, using HOMER^49^ [v4.8.3]. Each match must receive a sequence similarity score greater than or equal to the threshold provided by the PWMs (position weight matrices) or by HOMER (**Table S2**). We only scan for the TF motif of the corresponding TF ChIP-seq dataset, e.g. we only looked for T-BET motifs in the T-BET ChIP-seq data. We retained the highest scoring motif match for each ChIP-seq peak and used the genomic coordinates of the center two nucleotides to create a *GenomicRanges* object in R. For every run of IMPACT, 1,000 regulatory ranges are randomly selected, labeled with a 1, to train the model. Controlling the number of ranges used will standardize the logistic regression output such that predictions and model fits will be more comparable between cell-state models.

To build the non-regulatory class, we scanned the entire genome for TF motif matches, again using HOMER, and selected motif matches with no overlap with that TF’s ChIP-seq peaks, e.g. we scan for T-BET motifs, and only retain regions not overlapping T-BET ChIP-seq peaks. We do not check for overlap with other TF (i.e. GATA3, STAT3, FOXP3) ChIP-seq peaks. Similarly to the regulatory set, we used the genomic coordinates of the center two nucleotides of retained motifs to create the non-regulatory set, and label them as 0 in the regression. The motif matching process in both classes serves as a modest control for sequence content, as motifs are conserved regions of DNA. For every run of IMPACT, 10,000 regions of the non-regulatory set are randomly selected to train the model. This value is one order of magnitude larger than the regulatory set to reflect that genome-wide, we expect far more non-regulatory than regulatory elements. We justify setting this value to 10,000 if we assume that ~10% of the genome is regulatory, then for every positive region, we need nine negative regions. This would require 9*1,000 regulatory elements = 9,000 non-regulatory elements, a conservative estimate of the number of non-regulatory elements we actually use.

The sets of regulatory and non-regulatory elements are first partitioned into a random sampling of 80% each, to be used for 10-fold cross validation (CV), in which these sets are further partitioned into 90%/10% train/test. The remaining 20% to be used as a validation set (data completely unseen by the CV).

IMPACT is trained on standard 6-column bedfiles, of regions that are 2 base pairs wide, i.e.

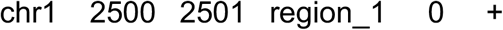

Epigenomic and sequence features are represented twice in the model, first with respect to local regions, and secondly with respect to distal regions. In the local case, for each region we annotate, we iterate through all genome-wide features, asking if there is overlap. In the distal case, we look for feature overlap of a more distal nucleotide (i.e. 1,000 base pairs up *or* downstream, such that overlap at either distal position will count; we do not distinguish between the upstream or downstream overlap in the feature matrix). Our rationale is that although a nucleotide may not intersect a particular feature, it may be informative to know that there is one nearby. The upstream and downstream distal coordinates (parameter set to 1,000 bp) are computed by subtracting or adding, respectively, the parameter value to these coordinates. If the computed distal coordinate is negative, the value is replaced with 1. If the computed distal coordinate is larger than the length of the chromosome, the value is replaced with the length of the chromosome. We set our distal parameter to 1,000 bp (**Table S3**) and do not check for overlap of any additional nucleotides within this distance. The feature matrix, on which the model is trained, may look like the following with dimensions (11,000 rows by 2+2X columns):

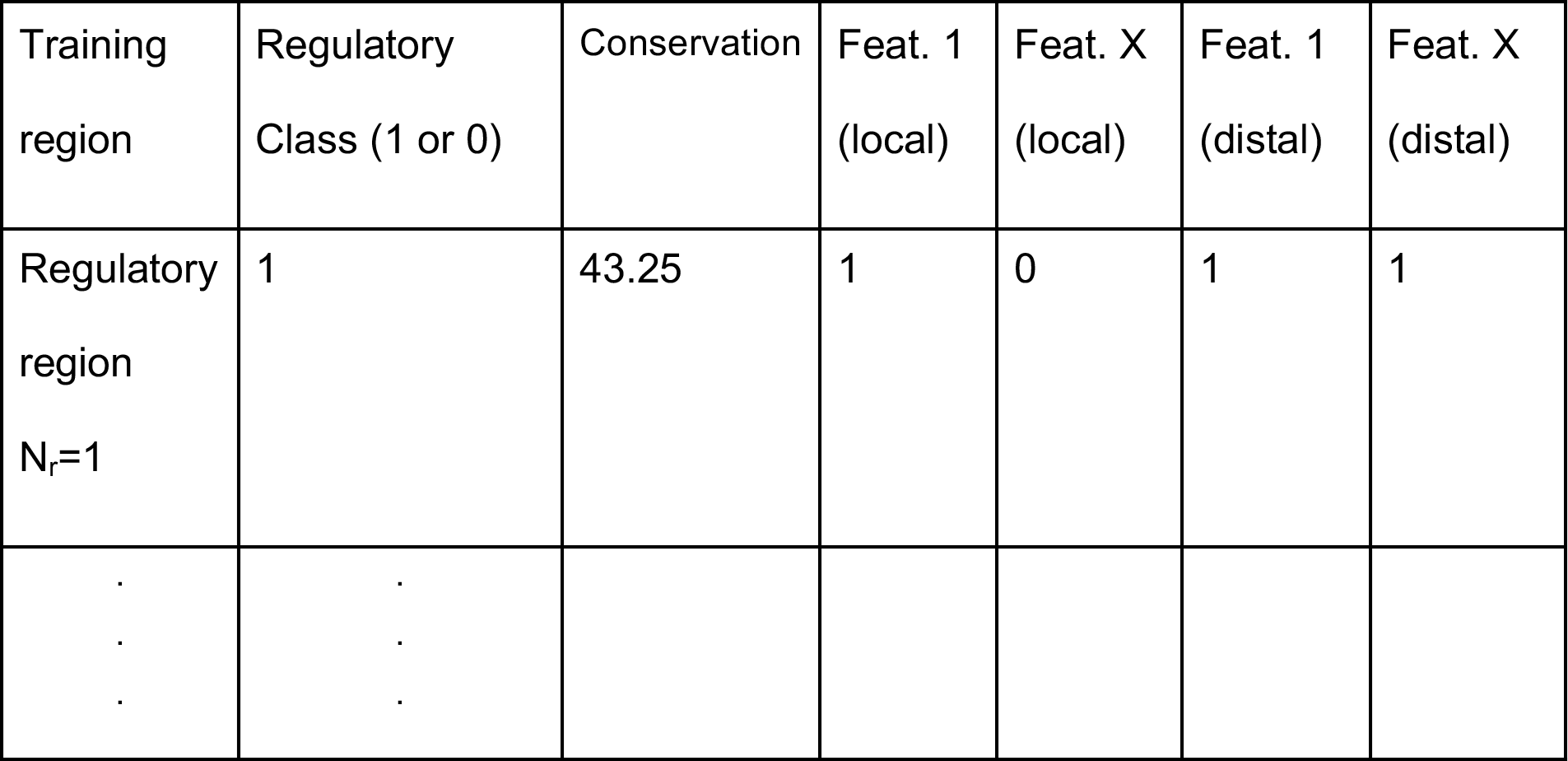

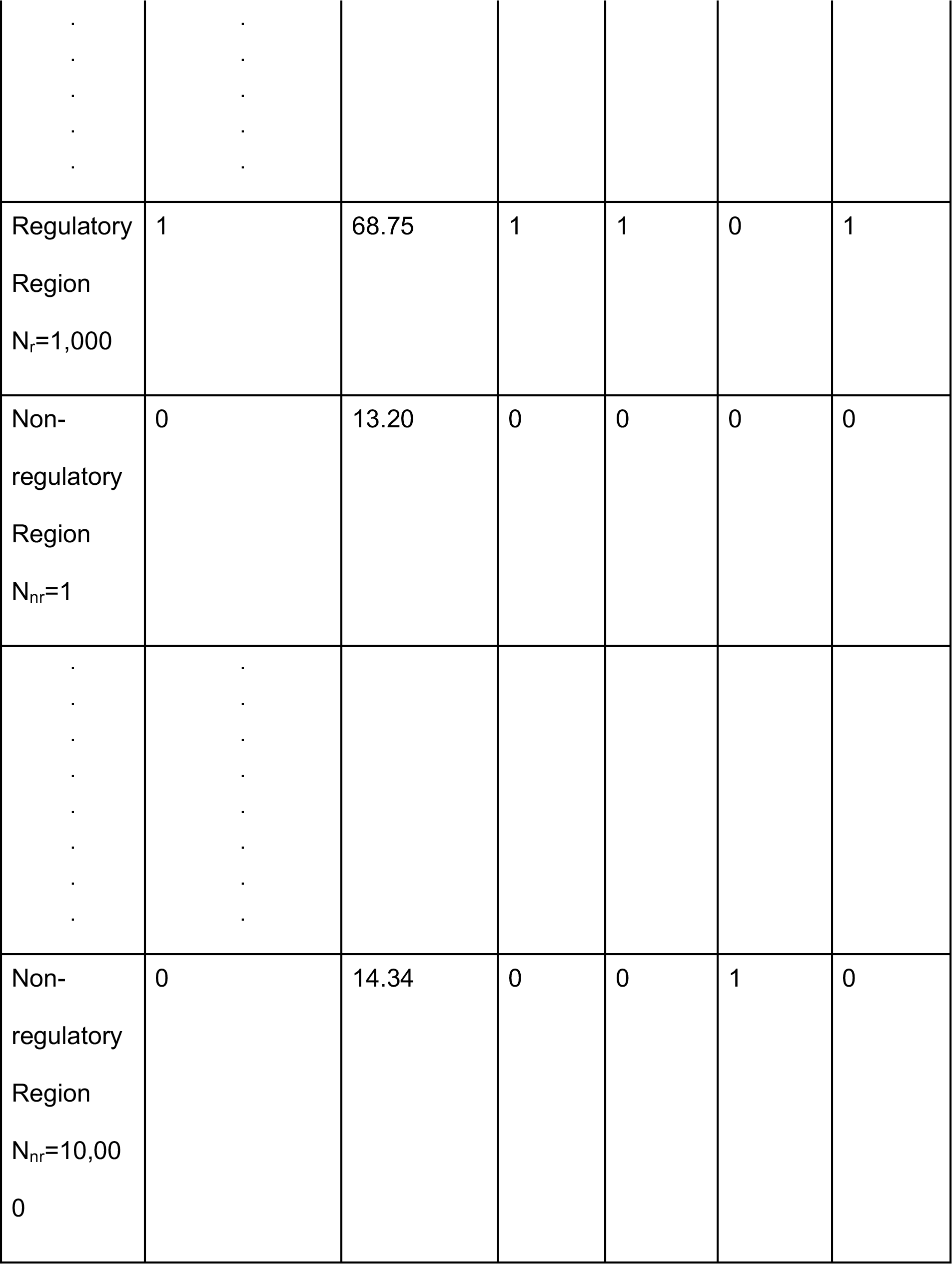

We applied IMPACT genome-wide to assign nucleotide-resolution regulatory probabilities, using the model *β* learned from the elastic net logistic regression CV. We show the largest magnitude *β* for each cell-state IMPACT model in **Figure S2**.

### Stratified-LD Score Regression (S-LDSC)

#### Genome-wide association data

We collected RA GWAS summary statistics^31^ for 38,242 European individuals, combined cases and controls, and 22,515 East Asian individuals, comprised of 4,873 RA cases and 17,642 controls^32^. Reference SNPs, used to estimate European LD scores, were the set of 9,997,231 SNPs with minor allele count greater or equal than five in a set of 659 European samples from phase 3 of 1000 Genomes Projects^50^. The regression coefficients were estimated using 1,125,060 HapMap3 SNPs and heritability was partitioned for the 5,961,159 reference SNPs with MAF ≥ 0.05. Reference SNPs, used to estimate East Asian LD scores, were the set of 8,768,561 SNPs with minor allele count greater or equal than five in a set of 105 East Asian samples from phase 3 of 1000 Genomes Projects^50^. The regression coefficients were estimated using 1,026,051 HapMap3 SNPs and heritability was partitioned for the 5,469,053 reference SNPs with MAF ≥ 0.05. Frequency and weight files (1000G EUR phase3, 1000G EAS phase3) are publicly available and may be found in our URLs.

#### Methodology

We apply S-LDSC^7^ [v1.0.0], a method developed to partition polygenic trait heritability by one or more functional annotations, to quantify the contribution of IMPACT cell-state-specific regulatory annotations to RA and other autoimmune disease heritability. We annotate common SNPs (MAF ≥ 0.05) with multiple cell-state-specific IMPACT models, assigning a regulatory element score to each variant. Then S-LDSC was run on the annotated SNPs to compute LD scores. Here, the two statistics we use to evaluate each annotation’s contribution to disease heritability are enrichment and standardized effect size (τ*).

If *a*_*cj*_ is the value of annotation *c* for SNP *j*, we assume the variance of the effect size of SNP *j* depends linearly on the contribution of each annotation *c*:

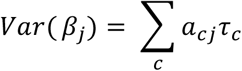

where τ_*c*_ is the per-SNP contribution from one unit of the annotation *a*_*c*_ to heritability. To estimate τ_*c*_, S-LDSC estimates the marginal effect size of SNP *j* in the sample from the chi-squared GWAS statistic 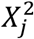:

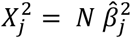

Considering the expectation of 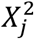 and following the derivation from Gazal et al 2017^9^,

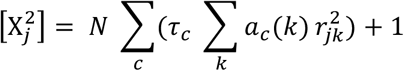

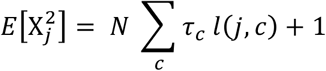

where *N* is the sample size of the GWAS, *l*(*j, c*) is the LD score of SNP *j* with respect to annotation *c*, and 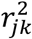 is the true, e.g. population-wide, genetic correlation of SNPs *j* and Since τ_*c*_ is not comparable between annotations or traits, Gazal et al 2017^9^ defines 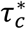 as the per-annotation standardized effect size, a function of the standard deviation of the annotation *c*, *sd*(*c*), the trait-specific SNP-heritability estimated by LDSC 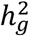, and the total number of reference common SNPs used to compute 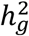, *M* = 5,961,159 in EUR and 5,469,053 in EAS:

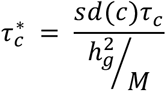

We define enrichment of an annotation as the proportion of heritability explained by the annotation divided by the average value of the annotation across the *M* common (MAF ≤ 0.05) SNPs. Enrichment may be computed for binary or continuous annotations according to the equation below, where 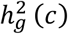 is the h2 explained by SNPs in annotation *c*.

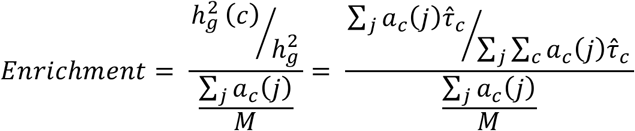

Enrichment does not quantify effects that are unique to a given annotation, whereas τ* does.

Each S-LDSC analysis involves conditioning IMPACT annotations on 69 baseline annotations, a subset of the 75 annotations referred to as the baseline-LD model. Our 69 annotations consist of 53 cell-type-nonspecific annotations^7^, which include histone marks and open chromatin, 10 MAF bins, and 6 LD-related annotations^9^ to assess if functional enrichment is cell-type-specific and to control for the effect of MAF and LD architecture. Consistent inclusion of MAF and LD associated annotations in the baseline model is a standard recommended practice of S-LDSC. When conditionally comparing two annotations, say A and B, in a single S-LDSC model, the two annotations may have similar enrichments if they are highly correlated. However, the τ* for the annotation with greater true causal variant membership will be larger and more statistically significant (e.g. > 0). Specifically, a τ* of 0, means that the annotation does not change per-SNP h2. A strongly negative τ ∗ means that membership to the categorical annotation decreases per-SNP h2, while a strongly positive τ* means that membership to the annotation increases per-SNP h2. The significance of τ* is computed based on a test of how different from 0 the τ ∗ is.

#### MHC exclusion

We note that S-LDSC excludes the MHC (major histocompatibility complex) due to its extremely high gene density and outlier LD structure, which is thought to be the strongest contributor to RA disease h2^51^. However, our work supports the notion that there is an undeniably large amount of RA h2 located outside of the MHC.

### eQTL enrichment

We acquired SNP-level summary statistics for 3,754 peripheral blood samples^23^. We computed a genome-wide enrichment of cis eQTL causal association across various functional annotations. To this end, we gathered summary statistics for each gene in the dataset, choosing only one probe when there was more than one tested on the array, and retained only the association statistics for the SNPs in a cis window of 1 Mb upstream and downstream of the gene TSS. For each gene in the context of each annotation, an enrichment was calculated explicitly as:

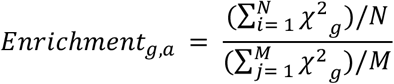

where *g* is the gene, *a* is the annotation, *N* is the number of variants within annotation *a*, *M* is the number of variants outside annotation *a*, *i* is the *i* ^*th*^ variant, *j* is the *j* ^*th*^ variant, and *χ*^2^ is the chi-squared statistic of the association between gene *g* and SNP *i* or *j*. We then computed genome-wide standard errors by block jackknifing the genome into 200 adjacent bins and computed a distribution of enrichment values when leaving one bin out at a time. This strategy is designed to prevent the genes of any one region of the genome from dominating the enrichment statistic. Furthermore, we used a permutation strategy to establish a null distribution. To this end, we randomly permuted the chi-squared associations in the cis-window of each gene 1000 times, while matching on 50 LD bins across the cis-window, and recomputed the enrichment with each of the functional annotations.

### Posterior probability enrichment

Previous work from our group aimed to define the most likely causal RA variant for each locus harboring a genome-wide significant variant^40^. To this end, posterior probabilities were computed with the approximate Bayesian factor (ABF), assuming one causal variant per locus. The posteriors were defined as:

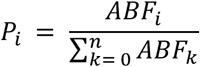

where *i* is the *i*^*th*^ variant, and *n* is the total number of variants in the locus. As such, the ABF over all variants in a locus sum to 1. Then, for each of the defined 20 RA-associated loci^40^, we computed the enrichment of high posterior probabilities in the top 1% of cell-state-specific IMPACT regulatory elements (**Table S5**). For each RA-associated locus *l*,

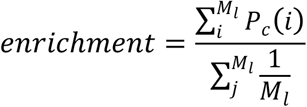

where *P*_*c*_(*i*) is the posterior causal probability of SNP *i*, such that *i* belongs to the top 1% of the cell-state-specific IMPACT annotation *c*, *M*_*l*_ is the number of SNPs in locus *l* for which we have a computed posterior probability. The denominator formulates the null hypothesis that each SNP in a locus is equally causal. We computed the average of these enrichment values over the 20 RA-associated loci. A permutation distribution was calculated by computing an average enrichment value over these 20 loci, in 1,000 trials, in which random posterior probabilities (of the same quantity *M*_*l*_) were selected. The permutation *p*-value was calculated by comparing the quantile value of the IMPACT enrichment to the assumed-normal permutation distribution defined by its mean and standard deviation, using the *pnorm* function in R.

### Caveats

This approach might be easily applied to a wide-range of other diseases, recognizing several important caveats. First, it relies on choosing a key regulator TF with a known consensus binding motif. Second, while for CD4+ T cells there is extensive literature of cell-states and their key regulatory drivers, this may not be readily available for all cell types. Third, it requires that primary cell ChIP-seq data, and therefore a specific antibody, are available for the desired TF in the disease-driving cell type. We prefer primary cell ChIP-seq data as opposed to cell line data in order to more closely approximate physiological regulatory element activity and chromatin dynamics. Fourth, as most immune populations are difficult to sort ex-vivo, collected cell-state-specific ChIP-seq data may not be 100% homogeneous. We expect that IMPACT predicted regulatory elements should be robust to experimental binding data of a heterogeneous population of cells. For instance, in a population of T cells that are 25% Tregs, and 75% other T lymphocytes, we expected more FOXP3 binding in the Tregs, as it is a specific regulator, relative to the other cells. Therefore, our TF occupancy profile should predominantly reflect that of Tregs. As there is especially poor availability of TF occupancy data in non-immune primary cells, we encourage the generation of more cell-state-specific data. Furthermore, due to IMPACT’s skew toward immune cell type features (~50%), we recommend supplementing relevant cell-type-specific features to avoid overfitting to immune cell types.

## Data Availability

All code for this paper has been made publicly available in the following GitHub repository: https://github.com/immunogenomics/IMPACT

## URLs

1. S-LDSC tutorial and instructions: github.com/bulik/ldsc
2. 1000G: www.1000genomes.org
3. RA EUR summary statistics: http://plaza.umin.ac.jp/yokada/datasource/software.htm
4. RA EAS summary statistics: http://jenger.riken.jp/en/result
5. 1000G Phase 3 LD scores, CD4+ T cell specifically expressed genes (binary functional annotations): http://data.broadinstitute.org/alkesgroup/LDSCORE/
6. Immgen.tsv: https://gist.github.com/nachocab/3d9f374e0ade031c475a

## Acknowledgements

We would like to thank Eric Lander and Jesse Engreitz for helpful discussions. This work is supported in part by funding from the National Institutes of Health (NHGRI T32 HG002295, UH2AR067677, 1U01HG009088, U01 HG009379 (Price/Raychaudhuri U01), and 1R01AR063759) and the Doris Duke Charitable Foundation Grant #2013097.

## Author Contributions

Developed IMPACT: T.A., Y.L., E.D.

Heritability analysis: T.A., Y.L., S.G., B.v.d.G. ATAC-seq: H-J.W., N.T.

RA genetic data: Y.O., K.Y., S.R.

RA disease/genetic interpretation: Y.O., K.Y., S.R. Statistical analysis: T.A., Y.L., S.G., B.v.d.G.

T.A. wrote the initial draft; all authors contributed to the final manuscript.

Work was conceived by T.A., S.R., A.P. and supervised by S.R. and A.P.

